# Efficient and reproducible generation of human iPSC-derived cardiomyocytes using a stirred bioreactor

**DOI:** 10.1101/2024.02.24.581789

**Authors:** Maksymilian Prondzynski, Raul H. Bortolin, Paul Berkson, Michael A. Trembley, Kevin Shani, Mason E. Sweat, Joshua Mayourian, Albert M. Cordoves, Nnaemeka J. Anyanwu, Yashasvi Tharani, Justin Cotton, Joseph B. Milosh, David Walker, Yan Zhang, Fujian Liu, Xujie Liu, Kevin K. Parker, Vassilios J. Bezzerides, William T. Pu

**Affiliations:** Department of Cardiology, Boston Children’s Hospital, Boston, MA 02115, USA; Disease Biophysics Group, Wyss Institute for Biologically Inspired Engineering, John A. Paulson School of Engineering and Applied Sciences; Fuwai Hospital, Chinese Academy of Medical Science, Shenzhen. Shenzhen, Guangdong Province, 518057, China; Harvard Stem Cell Institute, Cambridge, USA

## Abstract

In the last decade human iPSC-derived cardiomyocytes (hiPSC-CMs) proved to be valuable for cardiac disease modeling and cardiac regeneration, yet challenges with scale, quality, inter-batch consistency, and cryopreservation remain, reducing experimental reproducibility and limiting clinical translation. Here, we report a robust cardiac differentiation protocol that uses Wnt modulation and a stirred suspension bioreactor to produce on average 124 million hiPSC-CMs with >90% purity using a variety of hiPSC lines (19 differentiations; 10 iPSC lines). After controlled freeze and thaw, bioreactor-derived CMs (bCMs) showed high viability (>90%), interbatch reproducibility in cellular morphology, function, drug response and ventricular identity, which was further supported by single cell transcriptomes. bCMs on microcontact printed substrates revealed a higher degree of sarcomere maturation and viability during long-term culture compared to monolayer-derived CMs (mCMs). Moreover, functional investigation of bCMs in 3D engineered heart tissues showed earlier and stronger force production during long-term culture, and robust pacing capture up to 4 Hz when compared to mCMs. bCMs derived from this differentiation protocol will expand the applications of hiPSC-CMs by providing a reproducible, scalable, and resource efficient method to generate cardiac cells with well-characterized structural and functional properties superior to standard mCMs.

## Introduction

Numerous cardiac differentiation protocols have been established to differentiate human induced pluripotent stem cells (hiPSCs) cultured in adherent monolayer (ML)^1,2^ or three dimensional (3D) suspension^3–7^ formats. However, generation of high quantity and quality hiPSC-derived cardiomyocytes (hiPSC-CMs) with low functional variability between differentiations has remained challenging. Specific reasons include insufficient quality assessment of hiPSCs^8^, variability in cardiac differentiation outcomes^1,4^, and loss of hiPSC-CM functional properties following cryopreservation^9^. These and other limitations associated with low functional reproducibility (e.g., high cost and low translational value) motivate the development of improved cardiac differentiation protocols with defined benchmarks and consistent functional output from cryopreserved hiPSC-CMs.

Presently, most cardiac differentiation methods are based on programmed activation and then inhibition of the Wnt signaling pathway^2^ (Table S1). Due to its relative simplicity and high efficiency, this strategy became the dominant approach for hiPSC-CM differentiation for disease modeling and therapeutic cardiomyocyte replacement^10–14^. Typically performed in ML adherent cultures, this method has become the *de facto* standard for iPSC-CM differentiation. Nevertheless, hiPSC-CM differentiation by Wnt modulation in ML adherent culture has a number of important limitations. ML cultures scale poorly, with culture plate area and labor scaling linearly with cell number. Seeding density is a critical parameter for successful differentiation, and local heterogeneity in cell seeding reduces differentiation efficiency and increases well-to-well variation, causing low reproducibility in differentiation outcomes. Additionally, lack of flow within culture wells results in sub-optimal distribution of nutrients and pH buffering capacity.^1^ To improve hiPSC-CM purity, genetic reporter-, surface protein-, and lactate-based enrichment methods are often used following cardiomyocyte differentiation, but these purification methods can increase cost, create throughput barriers, or make hiPSC-CMs unsuitable for clinical applications^15^. While cryopreservation of cells is a critical step that dramatically facilitates design and execution of experiments and clinical translation, cryopreservation of ML-differentiated hiPSC-CMs negatively impacts their contraction, electrophysiology, and drug responses^9^. The limited scalability, well-to-well variation, and functional impact of cryopreservation combine to create significant barriers to the use of these cells.

To circumvent these limitations, suspension and stirred bioreactor cardiac differentiation protocols have been developed^3–7^. hiPSC-CMs obtained from suspension culture differentiation have been successfully utilized to model cardiac diseases^16–18^ or test novel molecular therapeutics^19^. Previous studies yielded 0.5-2 million hiPSC-CMs per mL for cultures ranging from 2.5 to 1000 ml^1,3,4^ (Table S1), illustrating the scalability of suspension cultures. However, different hiPSC lines exhibited inconsistent differentiation in previously reported suspension culture protocols^4^. The morphological, contractile, and electrophysiological properties of suspension and ML differentiated hiPSC-CMs has not been systematically evaluated, particularly after cryopreservation, although metabolic analyses suggested greater maturation of hiPSC-CMs from suspension cultures^20^.

Here we report an optimized bioreactor-based cardiac differentiation protocol with defined benchmarks enabling applicability to a variety of patient-, gene-edited and commercially available control iPSC lines without further modification and ensuring high quantity (∼124 million hiPSC-CMs per 100 ml run) and quality (>90% cTnT+ cells) of generated hiPSC-CMs. We describe protocols for robust cryopreservation and recovery of the bioreactor-derived hiPSC-CMs (referred to as bCMs) and assess their cellular composition via scRNAseq. We evaluated the morphological and functional properties of bCMs and compared them to ML-derived hiPSC-CMs (mCMs). bCMs had greater reproducibility and greater morphological and functional maturity.

## Results

### Optimized bioreactor differentiation protocol

We modified bioreactor and suspension cardiac differentiation protocols^4,5^ with goals of improving yield and reproducibility, increasing applicability across diverse iPSC lines, reducing cost, and enabling cryopreservation and recovery of resulting hiPSC-CMs. Our optimized workflow (Fig. 1a and S1a) built on a previously described embryoid body suspension culture protocols^4,5^ by incorporating the following features: (1) use of quality-controlled master cell banks (MCBs) to ensure consistency of input iPSCs; (2) use of a stirred bioreactor that continuously monitors and adjusts O2, CO2, and pH; (3) use of small molecules rather than growth factors, which are more expensive and vulnerable to lot-to-lot variation, to guide differentiation; (4) optimization of the time point to initiate differentiation by Wnt activation; (5) optimization of the duration of Wnt activation and the timing of Wnt inhibition; and (6) incorporation of controlled freeze and thaw protocols.

**Fig. 1:**
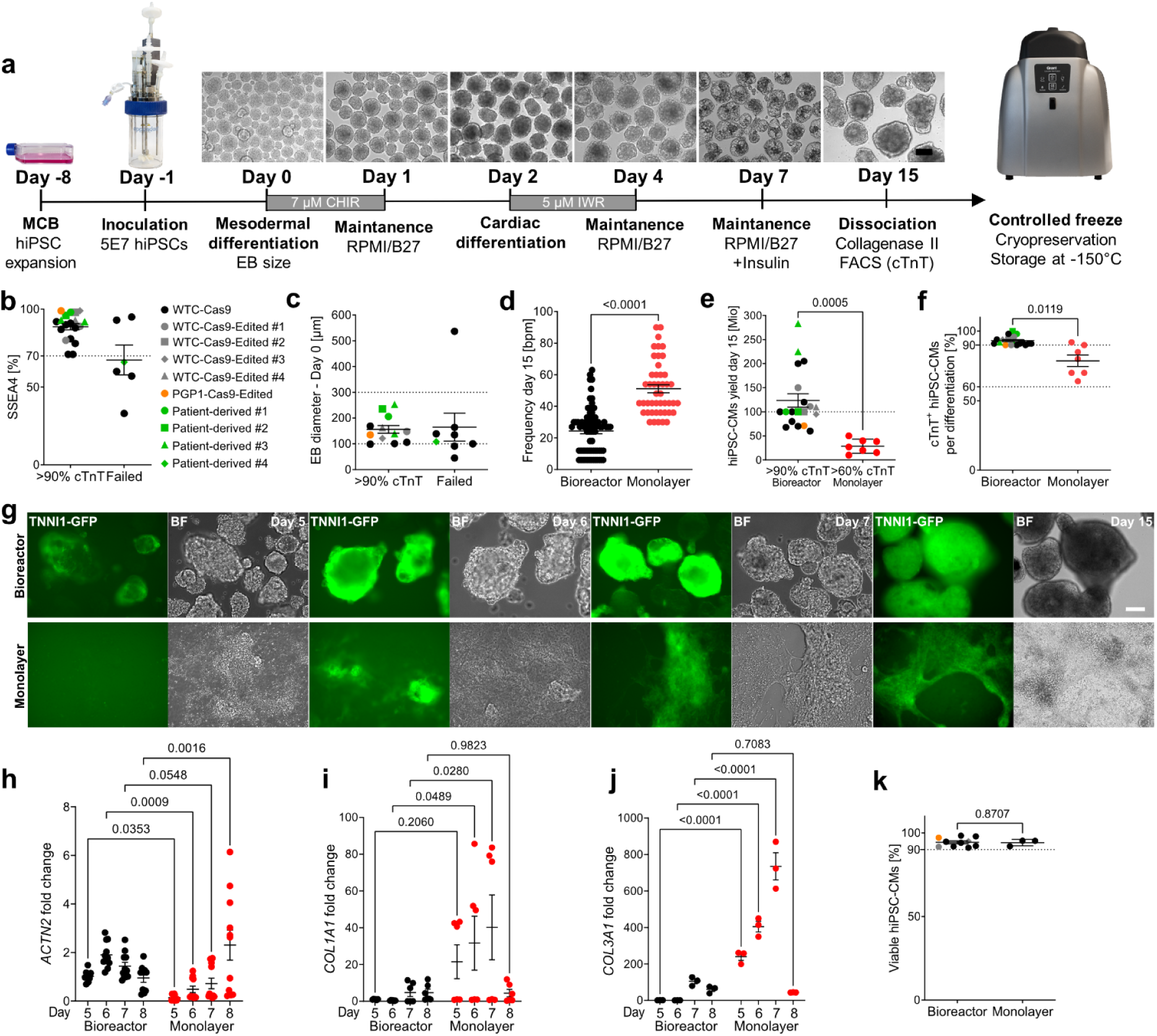
Optimized stirred bioreactor cardiac differentiation protocol. **(a)** Schematic representation of the optimized bioreactor cardiac differentiation protocol. **(b-c)** Characteristics of successful bioreactor cardiomyocyte differentiations. Runs were categorized as failed when they yielded < 90% hiPSC-CMs. hiPSC-CMs with frequency of pluripotency marker SSEA4 by flow cytometry (b) or out of range mean EB diameter (c) had higher likelihood of failure. n=19-25 differentiations. EB diameter analysis at day 0. **(d)** Spontaneous beating frequency of bioreactor- and monolayer-derived hiPSC-CMs at day 15 (n=number of differentiations; number of EBs or wells: Bioreactor (n=3; 71); Monolayer (n=3; 46). **(e-f)** hiPSC-CM yield (e) and purity (f) at day 15 of bioreactor or monolayer-directed cardiac differentiation (bioreactor, n=19 differentiations; monolayer, n=7 differentiations). Percentage of cells positive for cardiomyocyte marker cTnT was measured by flow cytometry. **(g)** Timeline of bioreactor and monolayer cardiac differentiation monitored by GFP fluorescence from a TNNI1-GFP iPSC line (bar: 200 µm). (**h-j**). qRT-PCR analysis of marker gene expression during bioreactor and monolayer cardiac differentiation. Values are expressed as fold-change compared to bioreactor day 5. *ACTN2* (**h**) marks cardiomyocytes, *COL1A1* (**i**) and *COL3A1* (**j**) mark fibroblasts. n=number of differentiations; number of replicates: Bioreactor (n=2-3; 6-11); Monolayer (n=2-3; 6-11). (**k**) Quantification of viable hiPSC-CMs after cryo-recovery from bioreactor (n=10) and monolayer (n=3) differentiations. Data are expressed as mean ± SEM. b-f,k, Welch’s unpaired *t*-test. h-j, two-way ANOVA with Holm-Šidák’s post-test. MCB, Master cell bank; human induced pluripotent stem cells,hiPSCs; EB, embryoid body; cardiac troponin T, cTnT; CM, cardiomyocyte; BF, bright field.

High quality input iPSCs are critical for successful and consistent differentiation^21^. Towards that end, we implemented procedures to establish MCBs of quality-controlled iPSCs (see Methods), including karyotyping (Fig. S1b) and mycoplasma testing. To monitor pluripotency of iPSCs input into the differentiation protocol, we measured pluripotency marker SSEA4 by FACS. High differentiation efficiencies (>90% expressing cardiomyocyte marker cardiac troponin T [cTnT]) were correlated with high SSEA4 (>70%) values, and low SSEA4 (<70%) predetermined failed differentiation (<90% cTnT+; Fig. 1b).

As in monolayer differentiation protocols^2^, we used Wnt activator CHIR99021 rather than growth factors to initiate mesoderm differentiation. We defined the optimal time of CHIR99021 addition based on EB diameter: EBs smaller than 100 μm fell apart upon CHIR incubation, and EBs bigger than 300 μm differentiated less efficiently (<90% cTnT+; Fig. 1c), likely due to inherent diffusion limits of larger EBs (Kempf et al. 2015; Manstein et al. 2021). Therefore our protocol targets CHIR99021 addition when EB diameter reaches 100 µm, which typically occurs at 24 hours.

We found that optimal cardiac differentiation occurs when CHIR exposure is limited to 24 hours, and when Wnt inhibition by addition of IWR-1 follows after an additional 24 hours (Fig. 1a and S1a). In 19 differentiations of 10 different iPSC lines treated with 7 μM CHIR and 5 μM IWR at these time points, we obtained on average ∼1.24 million cells per ml (Fig. 1e) with >90% cTnT+ cells (∼2.5 hiPSC-CMs/input hiPSC; Fig. 1f) over the course of 15 days. High percentages of cTnT positive cells were further confirmed by immunofluorescence analysis for cardiac markers cTnT and ACTN2 in cryosectioned bioreactor-derived cardiomyocytes (bCMs) at differentiation day 15 (Fig. S2c; Movie S1).

For comparison to bCMs, we used the same hiPSCs (WTC-Cas9) and differentiated them in parallel using the same differentiation protocol, except for a 48 h incubation period with CHIR instead of 24 h (Fig. S1a). Incubation with CHIR for 24 h led to failed ML differentiation. We did not apply hiPSC-CM enrichment methods^15^ at the completion of either bCM or mCM differentiation protocols. Compared to bCMs, ML-derived cardiomyocytes (mCMs) showed higher spontaneous beating frequency (Fig. 1d; Movie S2, S3), suggestive of lower maturity. Moreover, mCM differentiation resulted in lower cell yields (Fig. 1e) and higher variability in the fraction of cTnT+ cells per differentiation at day 15 (Fig. 1f).

We observed first contractions in bCMs at day 5 of differentiation (Movie S4), versus day 7 in mCMs and day 7 in previous reports of suspension culture hiPSC-CM differentiation^1,4–6^. To validate this observation, we differentiated hiPSCs in which sarcomere protein TNNI1 is fused to GFP^22^ in the bioreactor and visualized onset of GFP expression (Fig. 1g). TNNI1-GFP bCMs first expressed GFP at day 5 at the edges of spontaneously formed EBs, in regions that colocalized with areas of contraction. mCMs first showed GFP expression at day 6 (Fig. 1g). However, these GFP^+^ areas did not visibly contract. We also analyzed the time course of expression of the cardiac sarcomere gene *ACTN2*. By RT-qPCR, *ACTN2* was expressed in day 5 bCMs, whereas its expression level in mCMs did not become comparable until day 7. Additionally, we observed reduced inter-batch variation in *ACTN2* levels in bCMs compared to mCMs at days 7-15 (Fig. 1h and Fig. S2d). We also analyzed the expression of fibroblast markers *COL1A1* and *COL3A1*. These mRNAs were expressed at significantly lower levels in day 5 and 6 bCMs, suggesting a higher fraction of non-cardiomyocytes (non-CMs) in mCMs (Fig. 1i,j).

The ability to cryopreserve and recover viable cells that retain functional properties is critical to incorporate large scale differentiation protocols into efficient workflows. To enhance cryopreservation and functional recovery of hiPSC-CMs, we optimized cell dissociation protocols and cryo-protectant media, and achieved controlled rate cell freezing through a dedicated computer-controlled freezer. Our freeze/thaw protocol yielded ∼94% viable cells after recovery of bCMs and mCMs from cryopreservation (Fig. 1k). Functional testing of cryo-recovered hiPSC-CMs is discussed in subsequent sections.

Together, we established an integrated bioreactor-based workflow that yields consistently high numbers of highly pure hiPSC-CMs and developed methods for efficient cryo-preservation and cryo-recovery. The following sections further characterize the composition and functional properties of these cells.

### Cell composition of bCMs and mCMs

To gain a better understanding of generated cell types and differences between bCMs and mCMs, we performed single cell RNA sequencing (scRNAseq) of freshly dissociated hiPSC-CMs at day 15 of differentiation. Using microdroplet technology, we captured a total of 5,173 and 2,513 high quality single cell transcriptomes from bCMs or mCMs, each in biological duplicate. Unsupervised clustering on the most variable genes (Table S2) revealed 11 cell clusters and excellent agreement between biological duplicates (Fig. 2a; Fig. S3a). Based on expression of canonical marker genes, the clusters were identified as CMs (Clusters: 0, 1, 2, 4, 5, 8, 9), smooth muscle cells (Cluster: 6), non-CMs (Cluster: 3, 7), and endothelial cells (Cluster: 10) (Fig. 2b and S3b). The cardiomyocyte fraction was markedly higher in bioreactor (88%) compared to monolayer (51%) differentiation. We used canonical marker genes to classify the cardiomyocyte clusters. Clusters 0 and 1, highly enriched for ventricular marker genes *MYH7* and *MYL2*, were enriched in bioreactor (67% of cardiomyocytes) compared to monolayer differentiation (44% of cardiomyocytes; Fig. 2c). Conversely, cluster 2, containing cardiomyocytes with high expression of atrial marker genes *MYH6* and *MYL7*, were less frequent in bioreactor (8% of cardiomyocytes) compared to monolayer differentiation (36% of cardiomyocytes). Three cardiomyocyte clusters (4, 5 and 8) actively undergoing cell cycling (Fig. S3c) were present at comparable frequencies among cardiomyocytes (19% bioreactor vs. 16% monolayer). Compared to mCMs, bCMs expressed higher levels of mitochondrial metabolism genes *HADHA* and *ACADVL* (Fig. S3d,e), suggesting a higher degree of cardiac metabolic maturation, consistent with a prior report^20^. Upregulation of these genes was previously associated with a maturation protocol based on the induction of PPARdelta in hiPSC-CMs, which improved functional output^23^.

**Fig. 2:**
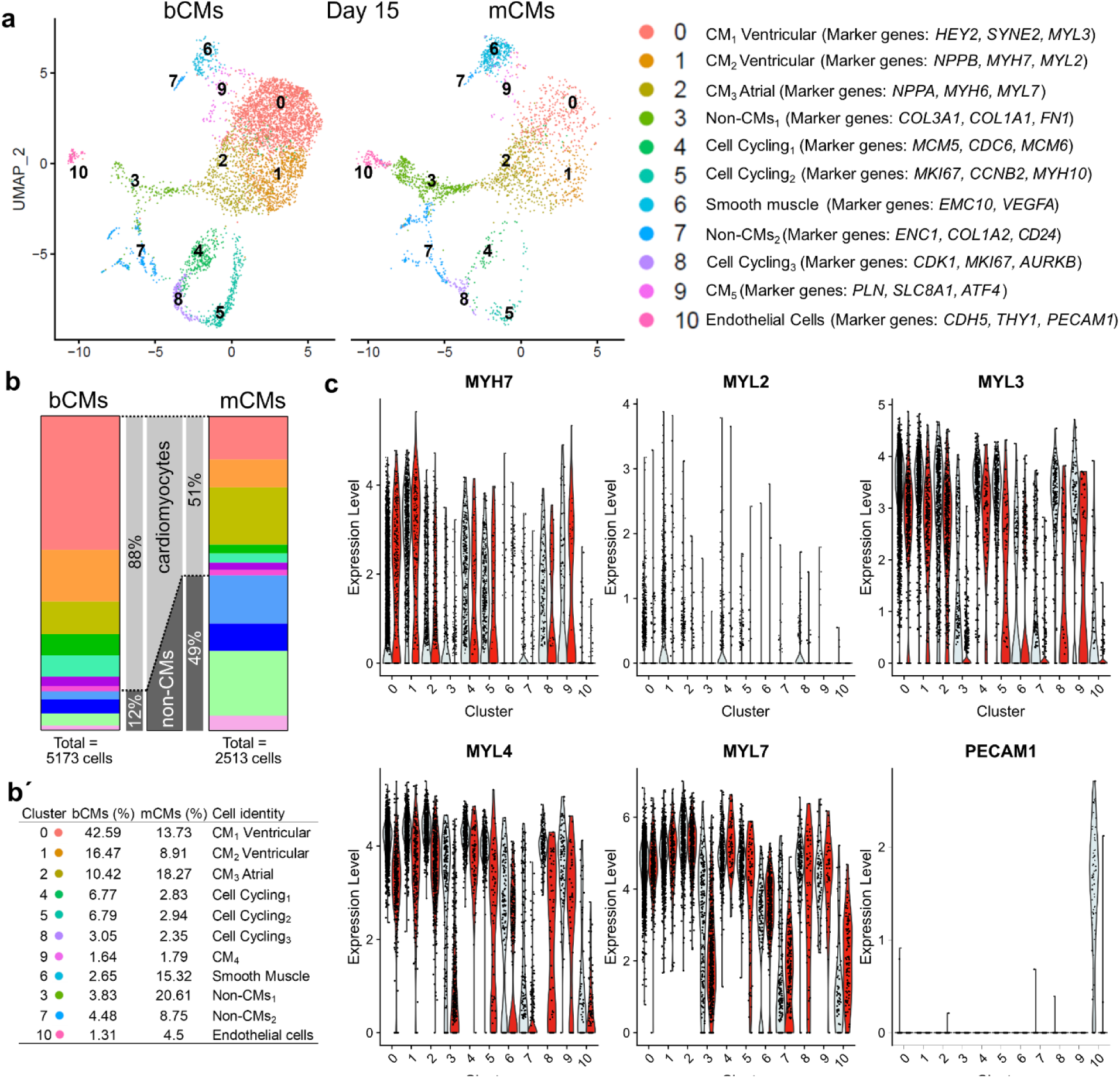
ScRNAseq reveals higher cardiomyocyte content and degree of cellular specification in bioreactor-derived hiPSC-CMs. **(a)** ScRNA-seq UMAP clustering of monolayer (ML) and bioreactor-derived hiPSC-CMs (bCMs) showing 11 clusters and their assigned cell types (right). **(b)** Stacked bar graph showing cellular composition of ML (left) and bCMs (right) expressed in % as parts of whole. Clusters are divided in cardiomyocytes (top) and non-CMs (bottom), whereby the color coding is the same as in panel A, and the order of clusters and the corresponding % are indicated in the table below (b**’**). **(c)** Violin-plot showing the relative expression of a subset of cardiac and non-cardiac marker genes (y-axis) across all clusters for bCMs (grey) and mCMs (red) (x-axis). Bioreactor-derived cardiomyocytes, bCMs; Monolayer-derived cardiomyocytes, mCMs; Noncardiomyocytes, non-CMs.

The non-cardiomyocyte fraction was dramatically lower in bioreactor compared to monolayer differentiation (12% vs. 49%; Fig. 2b, b’). An endothelial cell population marked by *PECAM1* was uniquely found in bioreactor differentiation (Fig. 2c). ML differentiation was highly enriched for non-cardiomyocyte clusters 3 and 7, which expressed a variety of marker genes for fibroblasts (*COL3A1*, *COL1A1*) and neurons (*CD24*, *ENC1*) suggesting a mixed or not fully established cellular identity in these cell populations (Fig. 2a, Fig. S3b, Table S2).

Taken together, scRNAseq analysis of cellular composition indicated that bioreactor differentiation yields a higher fraction of hiPSC-CMs, and these hiPSC-CMs have a greater degree of maturation and ventricular identity.

### Inter-batch reproducibility of bCMs in morphological and functional 2D assays

For morphological and functional analysis, cryopreserved hiPSC-CMs were thawed and plated at low density in 96-well plates pre-coated with diluted Geltrex (see Methods). After 7 days in culture, unpatterned hiPSC-CMs were fixed and morphologically analyzed. Staining for ACTN2 showed that many bCMs had elongated morphology, reminiscent of the rod shape of mature, *de facto* human cardiomyocytes. Quantification of bCM circularity confirmed their consistent elongated morphology across multiple batches. In comparison, mCMs displayed greater circularity and higher inter-batch morphological variation (Fig. 3b; Fig. S4a). bCM cell area was not significantly different than mCMs (bCMs: 1752 ± 112.3 μm^2^; mCMs: 1905 ± 246.6 μm; Fig. S4b), and inter-batch variation in cell area was significantly less than mCMs (Fig. 3c). Measured cell areas were comparable to other hiPSC-CM control lines cultured for 7^19^ and 30 days^16,18^. Mature, *de facto* human cardiomyocytes are 80% mononucleated^24^, and multinucleation tends to increase with cardiac disease^10,18,25^. Accordingly, both bCMs and mCMs were predominantly mononuclear. However, mCMs had elevated frequency of binucleated or multinucleated cardiac cells (Fig. 3d). Moreover, a greater fraction of mCM nuclei exhibited H2AFx immunoreactivity, a marker of DNA double strand breaks (Fig. 3e).

**Fig. 3:**
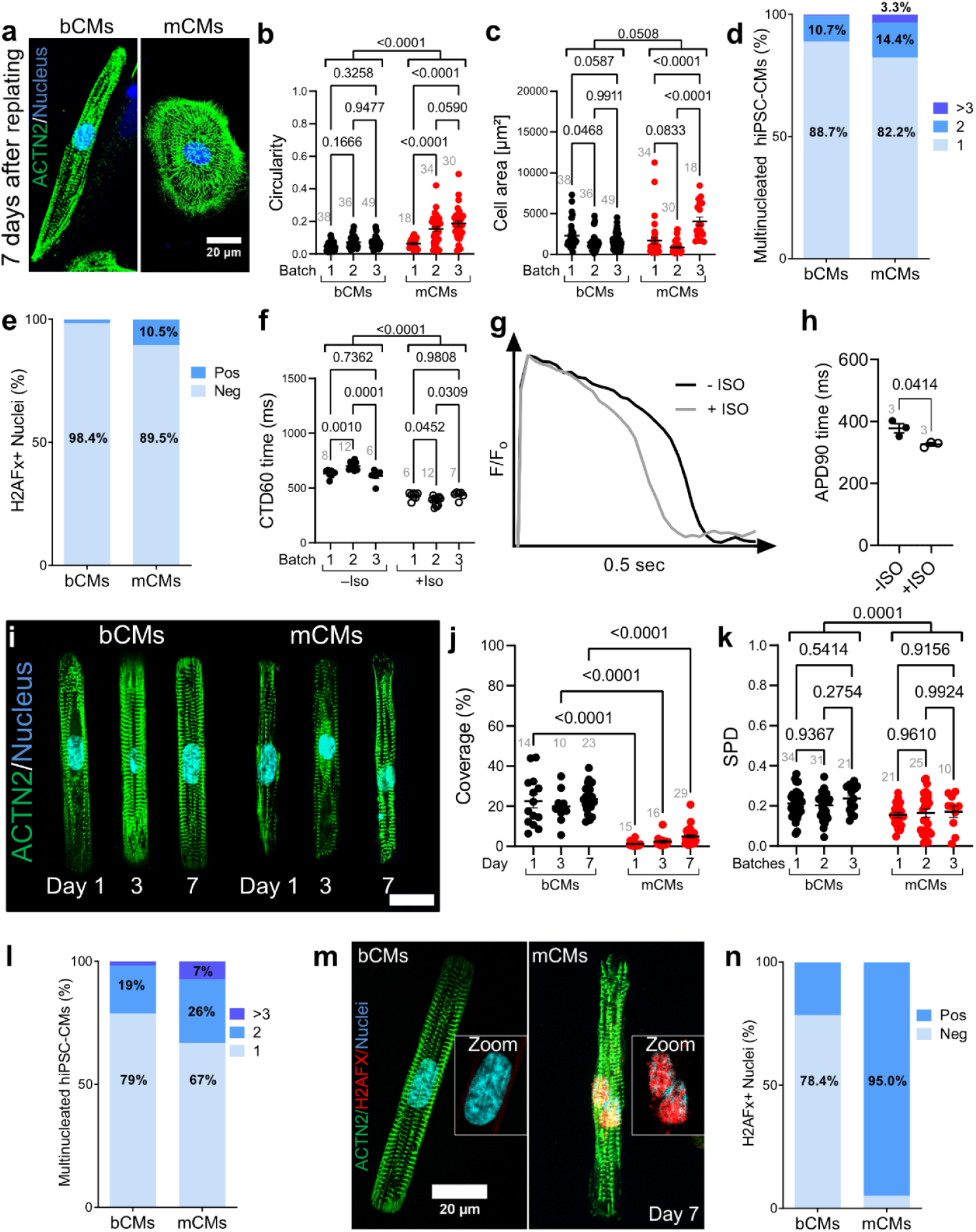
Comparison of bCMs and mCMs on 2D platforms. **(a-e)** Morphological characteristics of unpatterned hiPSC-CMs. Cryo-recovered bCMs and mCMs were cultured for 7 days on unpatterned Geltrex-coated dishes. Cells were stained for sarcomere Z-line marker ACTN2. Representative images (a) illustrate elongated shape of bCMs compared to mCMs. Bar, 20 µm. Circularity (b) and cell area (c) were quantified from 3 independent differentiation batches of bCMs and mCMs. Grey numbers indicate cells analyzed. Two-way ANOVA with Šidák’s post-test. **(d)** Nucleation of unpatterned bCMs (n=178) and mCMs (n=118) after 7 days in culture. Chi-squared p<0.0001. **(e)** Level of DNA damage response in unpatterned bCMs and mCMs. bCMs (n=63) and mCMs (n=95) were stained for H2AFX, which accumulates as a reaction to DNA double strand breaks. Chi-squared p<0.0001. **(f)** bCM calcium transients. Unpattered cryo-recovered bCMs were loaded with Ca^2+^ sensitive dye Fluo-4 and Ca^2+^ transients were optically recorded during 1 Hz pacing after 4 days of culture. mCMs could not be paced so were excluded. Calcium transient duration at 60% recovery (CTD60) was measured from 3 independent differentiation batches without (filled circles; n=26 wells) and with (open circles; n=13 wells) 1 µM isoproterenol (ISO) after 4 days in culture. Grey numbers indicate number of wells analyzed. Two-way ANOVA with Šidák’s post-test. **(g-h)** bCM action potentials. Unpattered cryo-recovered bCMs were loaded with voltage sensitive dye Fluovolt and optically recorded during 1 Hz pacing after 7 days in culture. g, Representative action potential traces for bCMs without and with (grey line) 1 µM isoproterenol (ISO). h, Quantification of action potential duration at 90% recovery (APD90) for bCMs without (n=3 wells) and with (n=3 wells) 1 µM isoproterenol (ISO). Three separate wells were analyzed per group. Data are expressed as mean ± SEM, paired t-test. **(i-k)** Characterization of micropatterned bCMs and mCMs. Cryo-recovered cells were plated on single cell extracellular matrix rectangular islands. Samples were fixed at stained after 1, 3, and 7 days in culture. i, Representative images. Bar, 20 µm. **(j)** Unbiased quantification of bCMs and mCMs coverage of single cell islands. Grey numbers indicate 10x fields analyzed. Two-way ANOVA with Šidák’s post-test. **(k)** Unbiased analysis of sarcomere packing density (SPD) of micropatterned bCMs and mCMs 7 days after replating. Grey numbers indicate cells analyzed. Two-way ANOVA with Šidák’s post-test. **(l)** Nucleation of micropatterned bCMs (n=192) and mCMs (n=60) after 7 days in culture. Chi-squared p<0.0001. **(m)** Level of DNA damage response in micropatterned bCMs and mCMs. Cells were stained for ACTN2 and H2AFX. m, representative images. Scale bar, 20 µm. **(n)** Quantification of H2AFx positive and negative nuclei in bCMs (n=51) and mCMs (n=20). Chi-squared p<0.0001. Data are expressed as mean ± SEM. Bioreactor-derived cardiomyocytes, bCMs; Monolayer-derived cardiomyocytes, mCMs; Isoproterenol, ISO.

Next, we analyzed the physiological properties of bCMs compared to mCMs. We recorded Ca^2+^ transients by loading hiPSC-CMs with the Ca^2+^ sensitive dye Fluo-4 and subjecting them to a 10 sec electrical pacing protocol. mCMs failed to follow electrical pacing (1 Hz; n=4 independent differentiation batches) and were therefore excluded from the analysis (Fig. S4c). In contrast, bCMs were reliably captured by the same pacing protocol. Ca^2+^ transients showed high inter-batch reproducibility and consistent responses to beta-adrenergic stimulation by isoproterenol (Iso) (Fig. 3f; Fig. S4d-h). Time-to-peak calcium transients in bCMs (218.6 ± 8.5 ms) were in agreement with a prior study using fresh and metabolically enriched mCMs derived from control line PGP1^26^. Interestingly, Hamad et al. reported on average 2-fold higher time-to-peak calcium transient times in control mCMs, which is comparable to the disease model studied by Psaras et al., suggesting a lower degree of maturation for mCMs that are not metabolically enriched^1,26^. Next, we recorded bCM action potentials (APs) by loading cells with Fluovolt, a membrane voltage sensitive dye, and pacing at 1 Hz. bCMs displayed typical ventricular AP morphology (Fig. 3g) 11 days earlier than previously reported for mCMs^1^, corroborating our scRNAseq findings at day 15 of differentiation (Fig. 2). As expected, adrenergic stimulation with Iso shortened action potential duration (Fig. 3g-h; Fig. S4i-j).

Together, these data indicate that bCMs possess low batch-to-batch variation and robust physiological and morphological cardiomyocyte properties, even after recovery from cryopreservation. In contrast, cryo-recovered mCMs had less mature morphology and could not be electrically paced.

### Micropatterned bCMs show a high degree of sarcomere maturation and survival

Plating hiPSC-CMs onto contact printed rectangular extracellular matrix (ECM) islands promotes their structural maturation, including alignment of sarcomeres perpendicular to the cell’s long axis^27^. We compared bCMs to mCMs after seeding onto rectangular ECM islands with the 7:1 aspect ratio of mature adult human cardiomyocytes. Cells were fixed on day 1, 3 and 7 after plating and stained for cardiac marker ACTN2 (Fig. 3l and S4k-k’). Cell attachment and sarcomere alignment were quantified by unbiased computational image analysis^28^. bCMs better survived plating on the micropatterned substrates, as demonstrated by their markedly higher coverage at all timepoints compared to mCMs (Fig. 3j; Fig. S4l). Sarcomere packing density and orientation order parameter, two different measures of sarcomere alignment with respect to the cardiomyocyte long axis, were considerably higher in bCMs than in mCMs at all investigated timepoints (Fig. 3k and Fig. S4m,n). Compared to unpatterned cells (Fig. 3d), a greater proportion of patterned bCMs and mCMs were bi- and multinucleated (Fig. 3l). Staining for DNA double strand break marker H2AFX indicated strikingly higher levels in patterned mCMs compared to patterned bCMs (Fig. 3m-n) or to unpatterned cells (Fig. 3e). Additionally, unbiased analysis of nuclear morphology^29^ identified a significantly higher fraction of nuclei with abnormal morphology in mCMs for all investigated timepoints (Fig. S5a-g). Together, these data indicate that bCMs are more amendable to single cell micropatterning than mCMs, with greater survival and sarcomere assembly, and reduced manifestations of genotoxic stress. Micropatterned substrates increased morphological maturation of bCMs, in line with previous findings on fresh mCMs^30,31^.

### bCMs show high force development and maturity in 3D engineered heart tissues

3D culture of hiPSC-CMs in fibrinogen gels subjected to anisotropic stress promotes cardiomyocyte maturation and sarcomere organization^5,16^. These engineered heart tissues (EHTs) also facilitate measurement of cardiomyocyte force development and relaxation. We assembled EHTs using cryopreserved bCMs (Movie S5) and mCMs (Movie S6; Fig. 4a). From days 5 to 32 after EHT casting, we recorded EHTs during spontaneous beating. bCM and mCM EHTs had similar spontaneous beating frequencies (Fig. 4b). bCM EHTs generated greater force than mCM EHTs at all investigated timepoints (Fig. 4c). Force generated by mCM EHTs (∼0.108 mN) were consistent with the prior reports for control hiPSC-CMs using the same EHT constructs^5,16^, whereas bCM EHTs force generation (∼0.384 mN) considerably exceeded these prior values. With increasing days in culture, bCM EHTs converged on similar contraction kinetics compared to mCM EHTs, as measured by the 50% contraction time (C50, Fig. 4d), and greater relaxation rates, as measured by the 50% and 90% relaxation time (R50, Fig. S6a; R90, Fig. 4e). Similarly, we recorded EHTs during optogenetic pacing at 1-3 Hz. At any given pacing frequency, bCM EHTs were captured more frequently than mCM EHTs (Fig. S6b). Under all pacing conditions, force was higher in bCMs EHTs compared to mCM EHTs (Fig. S6c), and C50 did not significantly differ between these groups (Fig. S6d). R90 was significantly lower in bCM EHTs at baseline and 2 Hz pacing; at 1 Hz and 3 Hz, bCM EHTs likewise had lower R90 values, although statistical significance was difficult to evaluate due to the low number of mCM EHTs captured at these rates (Fig. S6e). An independent set of EHTs was similarly analyzed with EHTs bathed in Tyrode solution, with similar findings (Fig. 4f-h; Fig. S6f). Under these conditions, both bCM and mCM EHTs generated greater force compared to culture medium, most likely due to higher calcium concentrations in Tyrode solution (bCMs: 0.384 mN vs 0.537 mN, mCMs: 0.108 mN vs 0.177 mN). Furthermore, in Tyrode solution a subset of bCM EHTs was successfully captured at 4 Hz pacing (Fig. S6g-j; Movie S8), notably faster than the previously reported maximal rates achieved for EHTs^32–34^ without physical conditioning^35,36^.

**Fig. 4:**
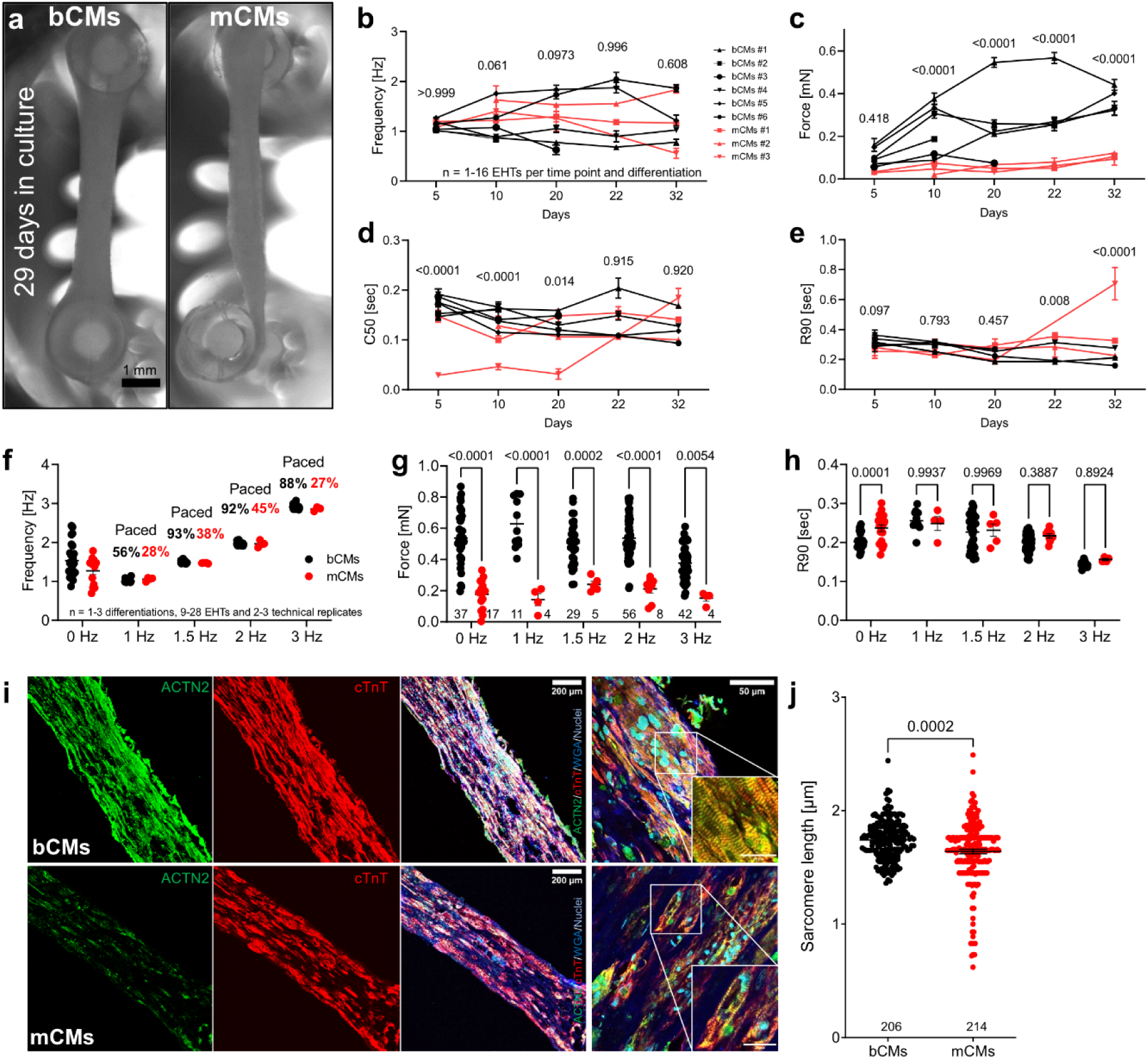
Comparison of 3D engineered heart tissues (EHTs) constructed with bCM or mCMs. **(a)** Representative images of EHTs after 29 days in culture (Scale bar, 1 mm). **(b-e)** Spontaneously beating EHTs assembled from cryo-recovered bCMs or mCMs were recorded in culture medium at 37°C from day 5 to day 32. Analyses of baseline frequency **(b)**, force **(c)**, time to 50% contraction (C50; **d**) and 90% relaxation (R90; **e**) showed greater force generation in bCM EHTs. Two-way ANOVA with Šidák’s post-test of pooled bCMs and mCMs values for each timepoint. **(f-h)** Analysis of EHTs in Tyrode solution (f-h) without pacing (0 Hz) or with 1-3 Hz pacing. **f**, EHT beat frequency in response to pacing. Only EHTs captured by pacing are shown. The percent of EHTs captured at each pacing rate is indicated. Two-way ANOVA with Šidák’s post-test. **(i)** Histological characterization of bCM and mCM EHTs. Cryosections of EHTs after 34 days in culture were stained for sarcomere Z-line protein ACTN2. Representative cryosections (i) showed higher cellularity and greater sarcomere content and organization in bCM EHTs. Sarcomere length (j) was quantified from 33 (bCM) or 35 (mCM) regions of interest from 2 (bCM) or 3 (mCM) EHTs. Data are expressed as mean ± SEM. Mann-Whitney test. Bioreactor-derived cardiomyocytes, bCMs; Monolayer-derived cardiomyocytes, mCMs.

Given greater force generation by bCMs in EHTs, we stained EHT cross-sections for cardiac sarcomere proteins ACTN2 and cTnT. Consistent with the physiological data, we found greater sarcomere density in bCM compared to mCM EHTs (Fig. 4i). Furthermore, bCM sarcomere length was higher in bCMs compared to mCMs EHTs (1.7 vs 1.6 µm; Fig. 4j), indicating greater maturity of bCMs. Overall, greater function of bCMs EHTs might be explained by the greater hiPSC-CM density, formation of more homogeneous tissue, and greater maturity including expression of metabolic genes *HADHA* and *ACADVL*, which have been previously associated with improved contractility in EHTs^23^.

## Discussion

Although cardiac differentiation protocols have been continuously improved over the past decade, current commonly used protocols suffer from limited scalability, high cost, inter-batch and even inter-well variation in efficiency and structural and functional properties, and loss of key functional properties upon recovery from cryopreservation^1,3–7,37^ (Table S1). These issues have presented substantial practical hurdles to performing reproducible and rigorous research using hiPSC-CMs. As a result, labs relying on these cells invest considerable resources in their continuous culture and differentiation and accounting for well and batch effects. Prior suspension culture and bioreactor protocols offered a potential solution with improved reproducibility and scalability, yet lacked robust methods for cryo-recovery, and the functional properties of the resulting cells were uncharacterized. Here we present a lower-cost bioreactor-based cardiac differentiation protocol (∼$569 per bioreactor run yielding ∼124 million hiPSC-CMs, compared to ∼$1075 for ML differentiation with comparable yield, exclusive of manpower; Table S3) with defined benchmarks and controlled freeze/thaw procedures. We extensively characterized the resulting bCMs and demonstrated that they have robust structural and functional properties of cardiomyocytes, reduced batch-to-batch variation, and greater force production in EHTs than previously described^5,16–18,32–34,36^. These levels of force were comparable to EHTs that underwent physical conditioning of increasing intensity^35^. Our hiPSC-CM bioreactor differentiation and cryopreservation pipeline and the subsequent deep morphological and functional characterization of bCMs provides a scalable and reproducible hiPSC-CM platform to enable subsequent disease modeling and therapeutic cardiomyocyte replacement.

We adapted the use of MCBs, small molecules, and optimized time points for Wnt activation and inhibition to increase efficiency in cardiac differentiation and reduced inter-batch variability. The success of this adaptation was reflected by first contractions observed as early as day 5 using our protocol (Movie S4), 2-3 days earlier than other protocols^1–5^. Spinner-^3^ and shaker-based differentiations^7^ reported higher yields of 1.5 to 2 million hiPSC-CMs per mL, but these approaches lack reproducibility in differentiation outcomes^3^, which were significantly improved by adding metabolic selection to the shaker-based protocol^7^. It has recently been shown that perfusion in the bioreactor enables high density hiPSC cultivation^38^. Our protocol might be further improved by applying perfusion and metabolic selection to increase yields and purities of generated hiPSC-CMs, respectively. Additionally, we optimized freeze/thaw procedures by utilizing a controlled rate freezer and serum-free freezing reagents^5,9^. All of these adaptations led to efficient cryo-recovery of bCMs with increased overall maturity and reproducible morphology, calcium transient durations and drug responses.

## Acknowledgment

The authors would like to thank Feng Xiao for providing hiPSCs for cardiac differentiation and Suellen Lopes Oliveira for assistance in graphic design.

## Funding Support

WTP and KKP were supported by the NCATS Tissue Chips Consortium (UH3 HL141798 and UH3 TR003279) and by charitable support from the Boston Children’s Heart Foundation. WTP and MP were supported by funding from Additional Ventures.

## Disclosures

The authors have no competing interests to disclose.

## Author contributions

M.P. and W.T.P. conceived the ideas and designed the experiments. M.P., R.H.B, P.B., M.A.T., K.S., M.E.S., J.M., A.C., N.J.A., Y.T., J.C., J.Mi., D.W., Y.Z., and X.L. conducted the experiments and analyzed the data. K.K.P. and V.J.B. contributed reagents and resources. M.P. and W.T.P. interpreted the data and wrote the manuscript.

## Supplemental Materials

### Online Methods

Data, analytic methods, and study materials will be made available on request to other researchers for purposes of reproducing the results or replicating the procedures. Studies were conducted under protocols approved by the Boston Children’s Hospital Institutional Review Board.

#### Human pluripotent stem cell culture

WTC-11 (Ctrl) is a wild-type human male iPSC line (Coriell Institute: # GM25256) that harbors a doxycycline (Dox)-inducible CRISPR/Cas9, which was created by introducing CAG-rtTA::TetO-Cas9 into the AAVS1 locus (Addgene #73500)^39^. The control iPSC line underwent cardiac differentiation in parallel in the bioreactor and in monolayer format followed by functional experiments and comparison to bioreactor-derived hiPSC-CMs was done using only the Ctrl line. Genome integrity was tested by digital karyotyping on the nanostring platform (Nanostring Technologies; (Fig. S1A). Additionally, cell culture supernatants were routinely screened for mycoplasma contamination using the LookOut® Mycoplasma PCR Detection Kit (Sigma, # MP0035-1KT) and master cell banks (MCBs) were established as described previously^21^.

Additional cell lines (RCM, Barth, Calmodulinopathy, ACM) were introduced into the Ctrl line using Cas9 genome editing, as described^40^. Patient-derived lines (1, 2, 3 and 4) were reprogrammed from blood using the CytoTune™-iPS 2.0 Sendai Reprogramming Kit (Thermo Fisher Scientific, # A16517).

#### Generation of hiPSC-CM in 2D monolayer culture

Monolayer cardiomyocyte differentiation was induced as previously described^2^ with some minor modifications (Fig. S1B). Briefly, Ctrl cells were washed once with PBS, detached by incubation with Versene (Invitrogen) for 5-10 min and seeded onto 12-well plates (Fisher Scientific # 0877229) pretreated with 1:100 (v/v) diluted Geltrex (Life Technologies, # A1413302) at a density of ∼45,000 cells/cm^2^ in E8 medium supplemented with 10 μM of ROCK inhibitor Y-27632 (R&D, # 1254). When cell confluency reached about 50-60%, cells were treated with basic medium (RPMI-1640 (# 61870127) plus 1x B27 minus insulin (# A1895601) all Life Technologies) containing 7 µM CHIR99021 (STEMCELL Technologies, # 72054) for 48 h, followed by basic medium containing 5 µM IWR-1-endo (STEMCELL Technologies, # 72564) for another 48 h. On day 7 of differentiation cells were cultured in basic medium supplemented with 1:1000 (v/v) insulin (Sigma-Aldrich, # I9278). Finally, cells were dissociated for 1-2 h depending on cell density using Collagenase II (Worthington, # LS004176) as described previously^5^ and below. Single cell suspensions were frozen in STEMdiff™ Cardiomyocyte Freezing Medium (STEMCELL Technologies, # 05030) using a controlled rate freezer (Grant, CRF-1) and stored in liquid nitrogen as described below.

#### Generation of hiPSC-CM in 3D bioreactor suspension culture

All bioreactor preparation steps were performed as described previously for the DASGIP® Parallel Bioreactor Systems (Eppendorf) with some minor modifications^4^. Inner vessel walls of the bioreactor (250 mL total volume) were coated with 1% (wt/vol) Pluronic F-127 (Sigma-Aldrich, # P2443) in PBS (Life Technologies, # 10010049) using a total volume of 50 mL for 1 h at 37°C. Coated vessels were washed three times with distilled water. In parallel, calibration of the pH sensor was performed at pH 4 (ACROS, # 61104-5000) and 7 (Fisher, # SB107-500) using commercially available buffer solutions. Upon completion all parts of the bioreactor were assembled (glass vessel, dissolved oxygen (DO) sensor, pH sensor, and head caps) and autoclaved at 121°C for 15 min. The sterilized bioreactor vessel was put under a laminar flow and 100 mL of PBS were added to each of the 250 mL volume capacity glass vessel (DASbox® Vessel, SR0250ODLS, 60 – 250 mL, overhead drive, 2 Rushton-type impellers, # 76SR0250ODLS). The vessel was placed onto the temperature controller and calibration of DO sensor was performed under agitation (60 rounds per minute; rpm) with 21% O_2_, 5% CO_2_ at 37°C and 10 standard liters per hour (sL/h) overlaying flow gas. After 15 days of cardiac differentiation (described below) the bioreactor set-up was disassembled and cleaned as described previously^4^.

Bioreactor cardiomyocyte differentiation was induced as previously described^2^ with modifications (Fig. S1C). At the start hiPSCs were cultured in T80 cell culture flasks (Life Technologies, # 178905) pretreated with 1:100 (v/v) diluted Geltrex (Life Technologies, # A1413302) incubated in a humidified incubator at 37°C, with 5% CO_2_ at a density of 15,000 cells/cm^2^. Cells were maintained in E8 medium (Life Technologies, # A1517001) with daily medium change until 90% cell confluency was reached. For dissociation, T80 flasks were washed once with PBS, incubated with 5 mL Versene (Life Technologies, # 1540066) for 15 to 20 min at 37°C and dissociation was stopped by adding 5 mL basic medium supplemented with 10 μM of ROCK inhibitor Y-27632. On average one flask resulted in 10 to 15 million hiPSCs depending on cell line and confluency upon dissociation. Cells were collected in 50 mL Falcon tubes and counted manually using a Neubauer Hemacytometer (Weber Scientific, # 3048-12). After centrifugation (200g, 5 min), 50 million single cells were resuspended in 50 mL E8 medium supplemented with 10 μM of ROCK inhibitor. The bioreactor vessel was taken from the bioreactor system and placed under the laminar flow. The PBS solution in the vessel was aspirated and replaced with 50 mL E8 (+ 10 μM Y-27632) and 50 mL of the hiPSC solution resulting in a final volume of 100 mL per vessel. Cells were agitated at a speed of 60 rpm, gassed with 21% O_2_ and 5% CO_2_ by 10 standard liters per hour (sL/h) overlay gassing and maintained at 37°C. The next day diameter of spontaneously formed embryoid bodies (EBs) was measured to estimate time of differentiation start. EB size was assessed by averaging at diameter measurements of at least 50 independent EBs. If critical diameter (100-300 µm) was reached, cardiac differentiation was induced by a complete change of the medium to RPMI 1640 with B27 (w/o Insulin; basic medium) supplemented with 7 μM CHIR99021 (day 0). After 24 h (Day 1), the complete medium was changed to basic medium and cells were incubated for an additional 24 h. On day 2, the complete medium was changed again to basic medium containing 5 µM IWR-1-endo for 48 h. If oxygen consumption was high and discoloration of the medium was visible on day 3, a 50% medium change using basic medium containing 5 µM IWR-1-endo was done. On day 4, the complete medium was changed to basic medium and cells were incubated for an additional 72 h with an optional 50% medium refreshment either on day 5 or 6. From day 7 on cells were cultured in basic medium supplemented with 1:1000 (v/v) insulin and 50% medium was refreshed on day 9, 11, 14. However, more medium refreshments were performed until day 15 if necessary. Finally, cells were dissociated on day 15 for 3-4 h depending on EB size and density, and frozen using a controlled rate freezer as described below.

#### Dissociation of hiPSC-CMs using Collagenase II

On day 15 monolayer and suspension differentiated hiPSC-CMs were dissociated using Collagenase II solution containing HBSS (Gibco, # 14175-095), Collagenase II (200 units per ml, Worthington, # LS004176), HEPES (10 mM, Sigma, # H3375), ROCK inhibitor Y-27632 (10 µM), and n-benzyl-p-toluenesulfonamide (BTS, 30 µM, VWR, # TCB3082)^5^. Monolayer differentiated hiPSC-CMs were washed twice with pre-warmed HBSS and incubated in Collagenase II solution (0.5 mL per 12-well) at 37°C in 5% CO_2_ for 1-2 h until single cells were apparent. For dissociation of suspension-derived hiPSC-CMs the bioreactor was taken from the temperature controller and opened under the laminar flow. The vessel was mixed manually to distribute all EBs equally in the vessel and 10 mL were taken out of the bioreactor and transferred into a 15 mL Falcon tube to estimate EB volume as described previously^5^.

Approximately, 12 mL of Collagenase II solution were used for 200 µL EB volume. Once EB volume was established, all EBs were pooled in a 50 mL Falcon tube and washed twice with HBSS, resuspended in Collagenase II solution, transferred to T175 suspension flasks (Sarstedt, # 83.3912.502) pretreated with 1% (wt/vol) Pluronic F-127, and incubated at 37°C in 5% CO_2_ for 3-4 h. Dissociation times vary, therefore, it is recommended to check after 1 h regularly on the progression by carefully tapping culture plates or flasks on a hard surface. If EBs were dispersing into single cells or very small clusters, they were gently triturated 5-10 times in their flask and transferred to a 50 mL Falcon tube. Dissociation reaction was stopped by adding the same volume of blocking buffer to the single cell suspension, which contained RPMI-1640 w/o B27 and DNase II (Sigma, # D8764), with 6 µL DNase II per mL RPMI-1640. Monolayer and suspension-differentiated hiPSC-CMs were counted, centrifuged (200g, 5 min) and prepared for downstream application or freezing (see below).

#### Freezing and thawing of hiPSC-CMs

For freezing, pelleted hiPSC-CMs were resuspended in cold (4°C) STEMdiff™ Cardiomyocyte Freezing Medium (STEMCELL Technologies, # 05030) and 1 mL was aliquoted into cryotubes (Fisher Scientific, # 12565163N) with cell numbers ranging from 1 to 20 million cells per cryotube. Cryotubes were frozen to -80°C in 1 h using a controlled rate freezer (Grant, CRF-1) followed by transfer to liquid nitrogen tanks (-150°C) for long-term storage.

For thawing of hiPSC-CMs a cryotube was removed from -150°C storage and immediately placed in a water bath set at 37°C. Constant moving of the cryotube in the water assured optimal distribution of heat and therefore uniform thawing of frozen cells. As soon as no ice crystal was visible cryotubes were placed under laminar flow and cell suspension was gently transferred to a 50 mL Falcon tube using a 1-ml pipette. The empty cryotube was rinsed with 1 mL of RPMI-1640 (room temperature) w/o B27 to recover residual cells, followed by dropwise addition to the 50-ml Falcon tube containing the cell suspension over 90 sec under gentle swirling. Next, 8 mL of RPMI-1640 (room temperature) w/o B27 were added to the cell suspension, whereby the first mL was added dropwise over 1 min and remaining 7 mL were added over 30 sec under gentle swirling. Finally, cell suspension was inverted three times in the 50 mL Falcon tube, counted using Trypan Blue (Sigma Aldrich, # T10282) and centrifuged at 200g for 5 min at room temperature. Pelleted hiPSC-CMs were resuspended and prepared for downstream applications as described below.

#### Flow cytometry

1×10^6^ single cells were added into a 15 mL Falcon tube, washed with 5 mL PBS, and centrifuged at 200*g* for 5 minutes. The supernatant was discarded and cells fixed for 10 min in 4% PFA, for intracellular epitopes, or ice-cold methanol (-20°C), for extracellular epitopes, followed by centrifugation at 200*g* for 5 minutes. Fixed cells were resuspended in 500 µL permeabilization buffer containing PBS, 5% (v/v) FCS/FBS (R&D systems, # S11150), 0.5% (w/v) Saponin (Sigma, # 47036-50G-F) and 0.05% (w/v) Sodium azide (Sigma, # S2002) and incubated overnight or 1 h at 4°C. Cell suspension was washed with 5 mL PBS, and centrifuged at 200*g* for 5 minutes. Pelleted cells were resuspended in permeabilization buffer containing directly labeled antibodies as shown in Table S4 and incubated for at least 45 min at 4°C in the dark. Cells were washed two times with 5 mL of PBS, the final pellet was resuspended in 200 μL PBS and cells analyzed by flow cytometry (LSR Fortessa Analyzer, BD Biosciences).

#### Gene expression

Total RNA from Cells, previously stored at −80°C, was extracted using an RNeasy Plus Universal Mini Kit (Qiagen, Valencia, CA, USA). The RNA integrity was assessed by automated electrophoresis using the RNA ScreenTape Analysis and 4200 TapeStation System (Agilent), concentrations were measured using a Nanodrop 8000 spectrophotometer (Thermo Scientific, Wilmington, DE, USA), and RNA was stored at −80°C. Synthesis of cDNA was performed with 0.5 μg total RNA using a High Capacity cDNA Reverse Transcription Kit (Applied Biosystems, Foster City, CA, USA), according to the manufacturer’s protocol in a MyCycler Thermal Cycler (Bio-Rad, Philadelphia, PA, USA). The cDNA was obtained in a final volume of 20 μL and stored at −20°C until it was used for the RT-qPCR expression assays. RT-qPCR was performed on the following genes using the SYBR green assay (Life Technologies # 4368708). PCR assays were carried out in 384-well plates using a CFX384 Touch Real-Time PCR Detection System (Bio-Rad). Relative expression was calculated using the 2-ΔΔCT method^41^, and results are presented as fold-change versus the control group mean values, normalized to GAPDH.

#### Single cell RNA sequencing

Collagenase II dissociated hiPSC-CMs on day 15 of cardiac differentiation and counted using a hemocytometer. Cells were resuspended in PBS + BSA (0.04%). Libraries were generated using the Chromium platform (10x Genomics) with the Next GEM Single Cell 3ʹ Reagent Kit v3.1, using an input of 1 million cells per mL. Gel-Bead in Emulsions (GEMs) were generated on the sample chip in the Chromium controller. Barcoded cDNA was extracted from the GEMs by Post-GEM RT-cleanup and amplified for 12 cycles. Amplified cDNA was then fragmented and subjected to end-repair, poly A-tailing, adapter ligation, and 10x-specific sample indexing following the manufacturer’s protocol. Libraries were quantified using the TapeStation (Agilent Technologies). Libraries were sequenced using NextSeq 500 (Illumina) at Harvard Medical School. Downstream differential expression and clustering analysis was performed using the Seurat V.4.0 package, as described in the tutorials (http://satijalab.org/seruat/). CellRanger matrices were imported for each sample with default parameters, and distributions of gene number, UMI number and % mitochondrial gene expression were examined for each sample to filter out cells of low quality. Cells with greater than 20% of genes coming from mitochondrial genes were selected against, as well as those with fewer than 200 genes or 400 UMIs. The doublets were detected and removed using DoubletFinder (v2.0.3). The resulting subset Seurat objects were normalized using the scTransform workflow and further scaled and normalized the RNA assay in order to perform downstream differential expression analysis and marker visualization utilizing the FindMarkers and FeaturePlot functions on the RNA assay. Uniform manifold approximation and projection (UMAP) was performed and iteratively modified after performing marker gene expression and examining expression of key markers. scRNAseq data will be made available upon publication of the revised manuscript.

#### Immunostaining of unpatterned and micropatterned hiPSC-CMs

hiPSC-CMs were thawed and cultured for 7 days in unpatterned 96-well plates (µclear®, Greiner Bio-One, 655090) and subsequently prepared for immunofluorescence analysis, as described previously^19^. The primary antibody ACTN2 (Sigma, A7811 (1:800), Table S4) was used followed by the secondary antibody anti-mouse Alexa Fluor® 488 (Life Technologies, LT A11029 (1:800), Table S4). Nuclei staining was obtained with Hoechst 33342 (5 µg/ml, Thermo Fisher Scientific). Images were obtained by confocal microscopy using a Zeiss LSM 800 or Olympus FV3000R confocal microscope with a 40x or 60x oil immersion objective.

Micropatterns were produced on glass coverslips (12 mm, VWR CAT NO 48366-252) coated with a 1:1 ratio of Polydimethylsiloxane 184 (10:1 Elastomer Base: Curing Agent, by The Dow Chemical Company LT H047M73001) and 527 (1:1 Part A : Part B, The Dow Company LT H047L6A013). Coverslips were coated for 46 hours in a 65°C oven. Then stamps with features of islands containing 7:1 aspect ratios^27^ were coated for 1 hour with Fibronectin (50 ug/ml) diluted in Geltrex (1:200, Life Technologies, # A1413302). In the meantime, PDMS-coated coverslips are exposed to UV Ozone for 8 minutes and the patterning process is performed by placing the dried stamps onto the coverslips. 1% Pluronics F-127 (Sigma-Aldrich, # P2443) is used to wash the coverslips for not more than 10 minutes, followed by washing 3 times with room-temperature PBS. To assess hiPSC-CMs in micropatterns 50,000 cells were plated on one micropatterned glass slide with a 7:1 size ratio and cultured for 1, 3 and 7 days. For immunofluorescence analysis cells were fixed with 4% PFA for 10 min at 4°C, followed by permeabilization in 0.1% Triton X-100 (Sigma-Aldrich, LT 069K0049) dissolved in PBS for 10 min at RT. hiPSC-CMs were incubated in blocking buffer (PBS, 3% BSA (Sigma, LT SLBW9820)) for 1 hour at room temperature and then incubated with conjugated antibodies for ɑ-actinin/FITC (1:50, Table S4) and Phalloidin 647 (1:400, Table S4) in addition to Hoechst (1:500) in staining buffer (PBS, 1% BSA (Sigma, LT SLBW9820)) for 2 hours at RT. Micropatterned glass slides were then mounted using Prolong Diamond Antifade Mountant (Invitrogen, LT 2273639). Images were obtained by confocal microscopy using a Olympus FV3000R confocal microscope with a 60x oil immersion objective.

#### Morphological analysis of hiPSC-CMs

Quantification of myofibrillar disarray and cell area was evaluated with Fiji (ImageJ) as described previously^16,19^. Additionally, values for circularity were captured with Fiji (ImageJ) by assigning a cell and using the “Measure” function. To assess sarcomere alignment in micropatterned hiPSC-CMs we used previously reported methods from the Disease Biophysics Group(Pasqualini et al. 2015) that utilizes ImageJ Plugins (OrientationJ) and a custom-made Matlab script for structural analysis of the cells Resulting parameters for sarcomere alignment referred to as “Orientational Order Parameter” (OOP) were captured, whereby the OOP2 value corresponded to the orthogonal alignment of actin bundles representing Z-disc regularity. Multinucleation was assessed by manually counting the number of nuclei per cell in unpatterned and micropatterned formats. Morphology of nuclei was assessed using a previously published Fiji plug in^29^.

#### Ca^2+^ and voltage imaging

For high throughput Ca^2+^ and voltage imaging, we used the Vala Biosciences Kinetic Image Cytometer (KIC). Cryopreserved hiPSC-CMs were thawed and seeded onto polystyrene 96 well plates (Greiner Bio-One, 655090) at 50,000 to 100,000 cells per well. After four days in culture, cells were stained with 0.1 µg/ml Hoechst 33342 (Life Technologies, cat) and 2.5μM Fluo-4 calcium indicator dye (Invitrogen F14201) or 1:1000 FluoVolt membrane potential kit (Invitrogen F10488) in Tyrode’s solution for 30 min at 37°C. Cells were then washed with Tyrode’s solution and imaged by KIC with a 20x objective at 67 frames per second. The imaging protocol consisted of 1Hz electrical stimulation with 10 seconds of pre-pacing followed by 10 seconds of video acquisition. Raw data was analyzed with CyteSeer software (Vala Biosciences) using custom scripts for analysis of calcium transient kinetics and action potential duration.

#### Generation of engineered heart tissues

Engineered heart tissues (EHTs) were generated as described previously^5,42^ with some minor modifications. Briefly, 0.8x10^6^ hiPSC-CMs were used to generate each EHT. Cells were transduced with adenovirus ChR2-YFP on the day of casting or after 7 days in vitro. We modified the standard EHT culture medium (EHT-medium in the referenced literature^5^) by replacing DMEM with RPMI 1640 plus B27 minus insulin, removing 10% heat-inactivated horse serum, and reducing aprotinin concentration to 5 µg/ml. This resulted in EHTs initiating contraction as early as day 1-3 after EHT assembly, versus day 7-10 reported in the literature^5^. EHT contraction was recorded as described below from day 7 on and functional analysis was performed from day 27 to day 33. For Immunohistochemical analysis EHTs were cryosectioned and stained with primary antibodies shown in Table S4. Sections were imaged using an Olympus FV-3000 confocal microscope and analyzed with Fiji (ImageJ).

#### Functional assessment of EHTs

EHTs in a 24 well plate were placed in a stage top incubator and maintained at 37C, 5% CO2. EHTs were optically paced at different frequencies using blue LEDs positioned above the place, and recorded from below at 30 frames per second through a 561 nm long-pass filter (Semrock BLP02-561R-32) using an 8mm f/1.4 lens (ThorLabs MVL8M1) mounted on a Basler acA1920 camera. EHT post movement was tracked post-hoc using the multi-template matching FIJI plugin^43^. Twitch force measurements were subsequently measured by applying post deflection to the beam bending theory for a known Young’s modulus of the posts, as described in detail elsewhere^44^.

**Fig. S1:**
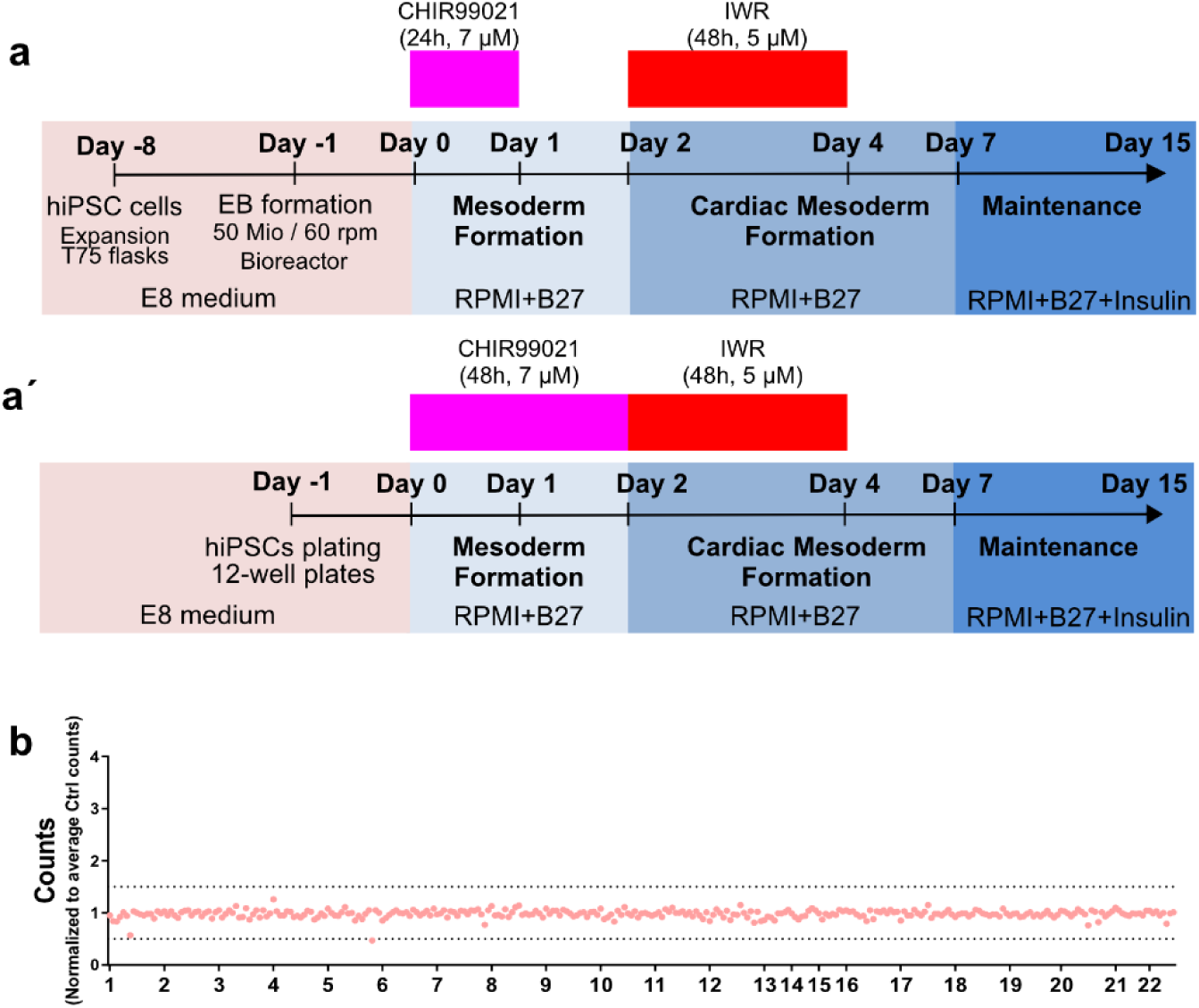
Cardiac differentiation protocols and iPSC karyotyping. Schematic representation of the cardiac differentiation protocol in suspension **(a)** and monolayer cultures **(á)**. **(b)** Digital karyotyping results of WTC control iPSCs at passage 56.

**Fig. S2:**
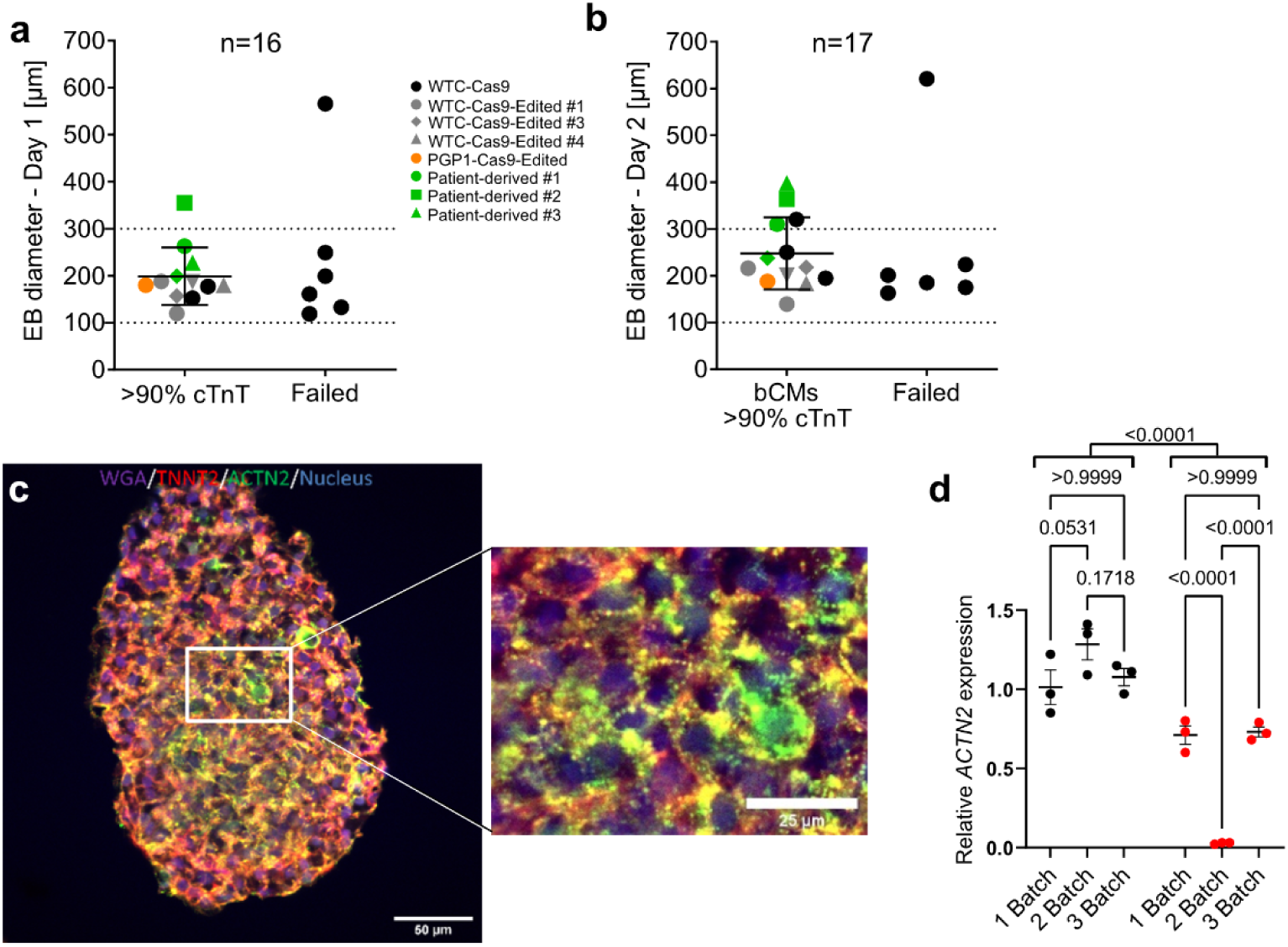
Characterization of embryoid bodies (EBs) generated with our optimized cardiac differentiation protocol. **(a-b)** EB diameter at day 1 (a) or 2 (b) and relationship to bioreactor differentiation outcome. Outcomes were classified as cTnT+ cells > 90% or ≤ 90% (“Failed”). **(c)** Representative cryosection of an EB at day 15 stained for wheat germ agglutinin (WGA), cardiac troponin T (cTnT), alpha actinin 2 (ACTN2) and Hoechst 33342. Boxed region is enlarged at right. Scale bar: 50 µm and 25 µm (zoom)). **(d)** qRT-PCR analysis of cardiac marker gene *ACTN2* at day 15 of cardiac differentiation. Points represent technical replicates for each independent differentiation. Two-way ANOVA with Bonferronís post-test. Data are expressed as mean ± SEM.

**Fig. S3:**
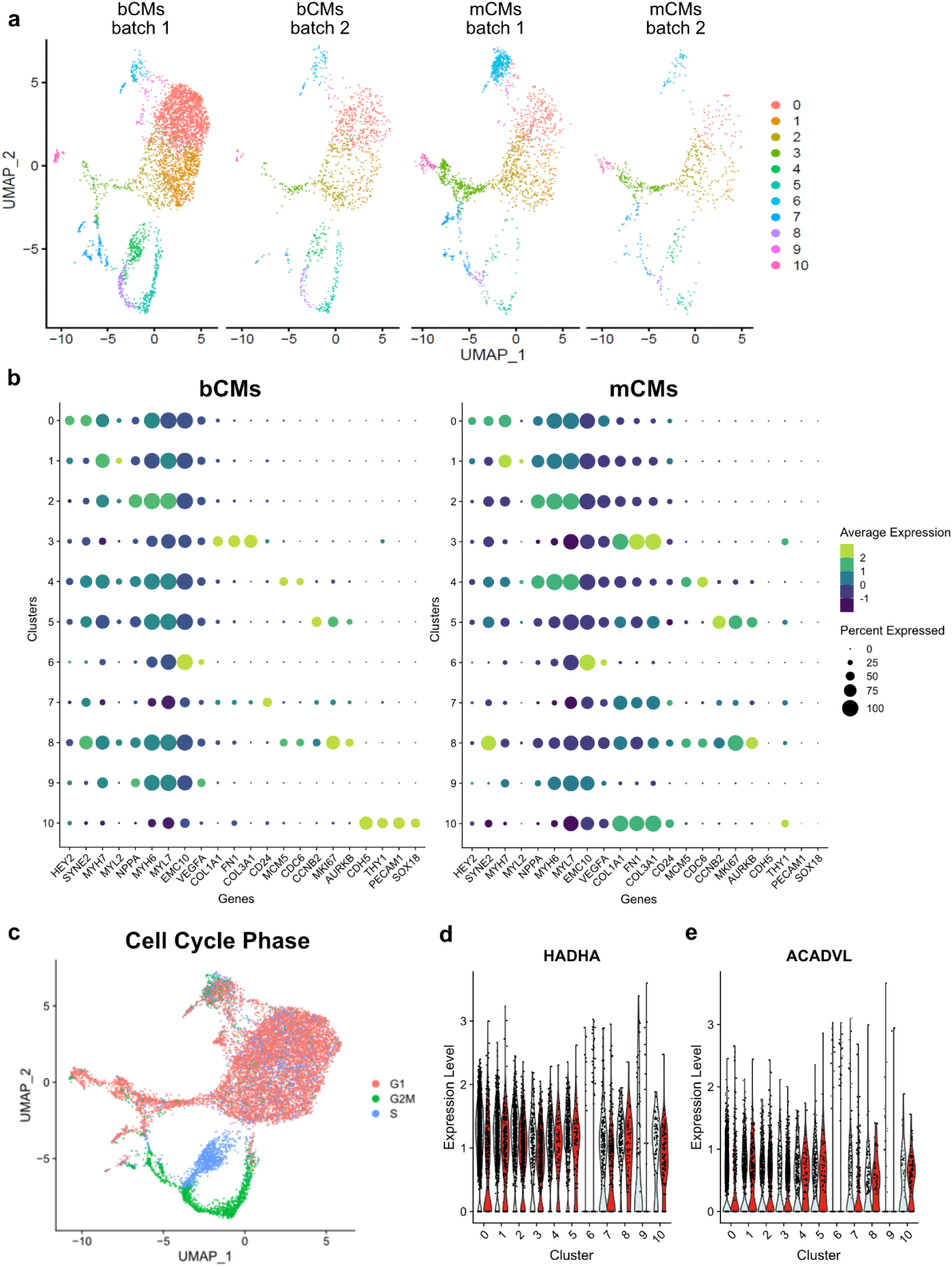
scRNAseq results for bioreactor and monolayer-derived hiPSC-CMs. **(a)** scRNA-seq UMAP clustering of mCMs (right) and bCMs (left) showing 2 biological replicates and corresponding cluster assignment. **(b)** Dot-plot showing the relative expression of a subset of cardiac and non-cardiac marker genes (x-axis) across all clusters (y-axis) for bCMs (left) and mCMs (right). **(c)** scRNA-seq UMAP plot showing the cell cycle stages G1, G2M and S phase for mCMs and bCMs. Violin-plot showing the relative expression of *HADHA* **(d)** and *ACADVL* **(e)** across all clusters for bCMs (grey) and mCMs (red). Bioreactor-derived cardiomyocytes, bCMs; Monolayer-derived cardiomyocytes, mCMs.

**Fig. S4:**
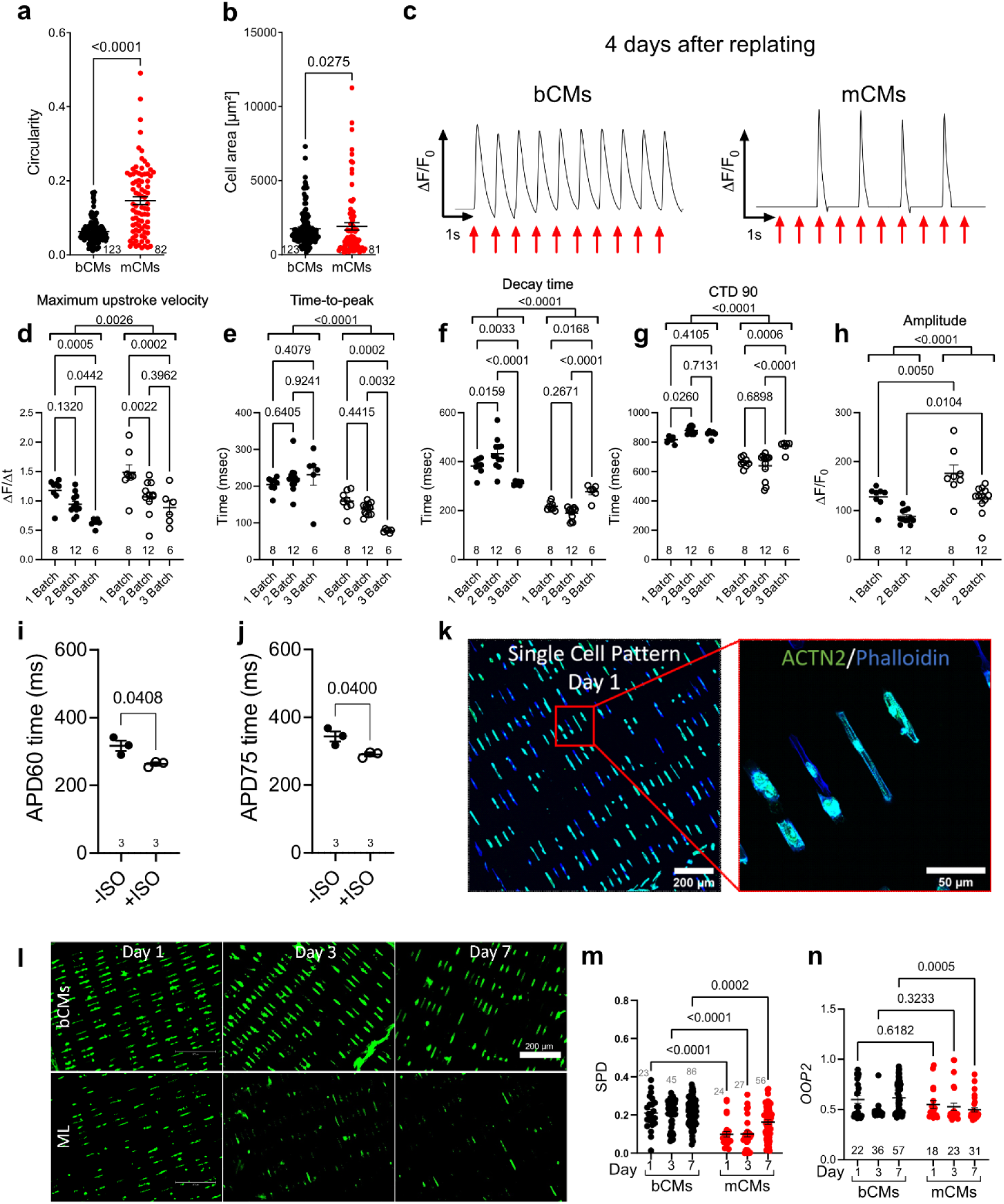
2D modeling of bCMs and mCMs. Circularity **(a)** and cell area **(b)** measurements of 3 independent differentiation batches for bCMs (n=123) and mCMs (n=82). Mann-Whitney test. **(c)** Representative calcium transients of bCMs (top) and mCMs (bottom) paced at 1 Hz. Pacing signals are indicated by the red arrows. (**d-h**) Calcium transients of Fluo-4 loaded bCMs were recorded with 1 Hz pacing. Maximum upstroke velocity **(d)**, time-to-peak **(e)**, decay time **(f)**, calcium transient duration (CTD) 90 **(g)** and amplitude **(h)** in the presence or absence of 1 µM isoproterenol (ISO) were compared using Two-way ANOVA with Šidák’s post-test. Number of wells quantified is indicated along the bottom of the plot. (**i-j**) Action potentials were recorded from Fluovolt-loaded bCMs paced at 1 Hz. Action potential duration (APD) 60 **(i)** and 75 **(j)** for bCMs in the presence or absence of 1 µM ISO were compared using paired *t*-test. **(k)** Representative image of bCMs fixed after 1 day in culture on micropatterned substrates and immunostained for ACTN2 and F-actin. Scale bar, 200 µm (k) or 50 µm (k’). **(l)** Representative images used to quantify coverage micropatterns by bCMs (top) and mCMs (bottom). Cells were immunostained for ACTN2. Scale bar, 200 µm. Unbiased quantification of sarcomere alignment. Computational image analysis calculated SPD **(m)** and OOP2 **(n)**, measures of sarcomere alignment, on bCMs and mCMs cultured on micropatterned substrates. Number of cells quantified is indicated along the bottom of the plot. Two-way ANOVA with Šidák’s post-test. Data are expressed as mean ± SEM. Bioreactor-derived cardiomyocytes, bCMs; Monolayer-derived cardiomyocytes, mCMs; ADP, action potential duration.

**Fig. S5:**
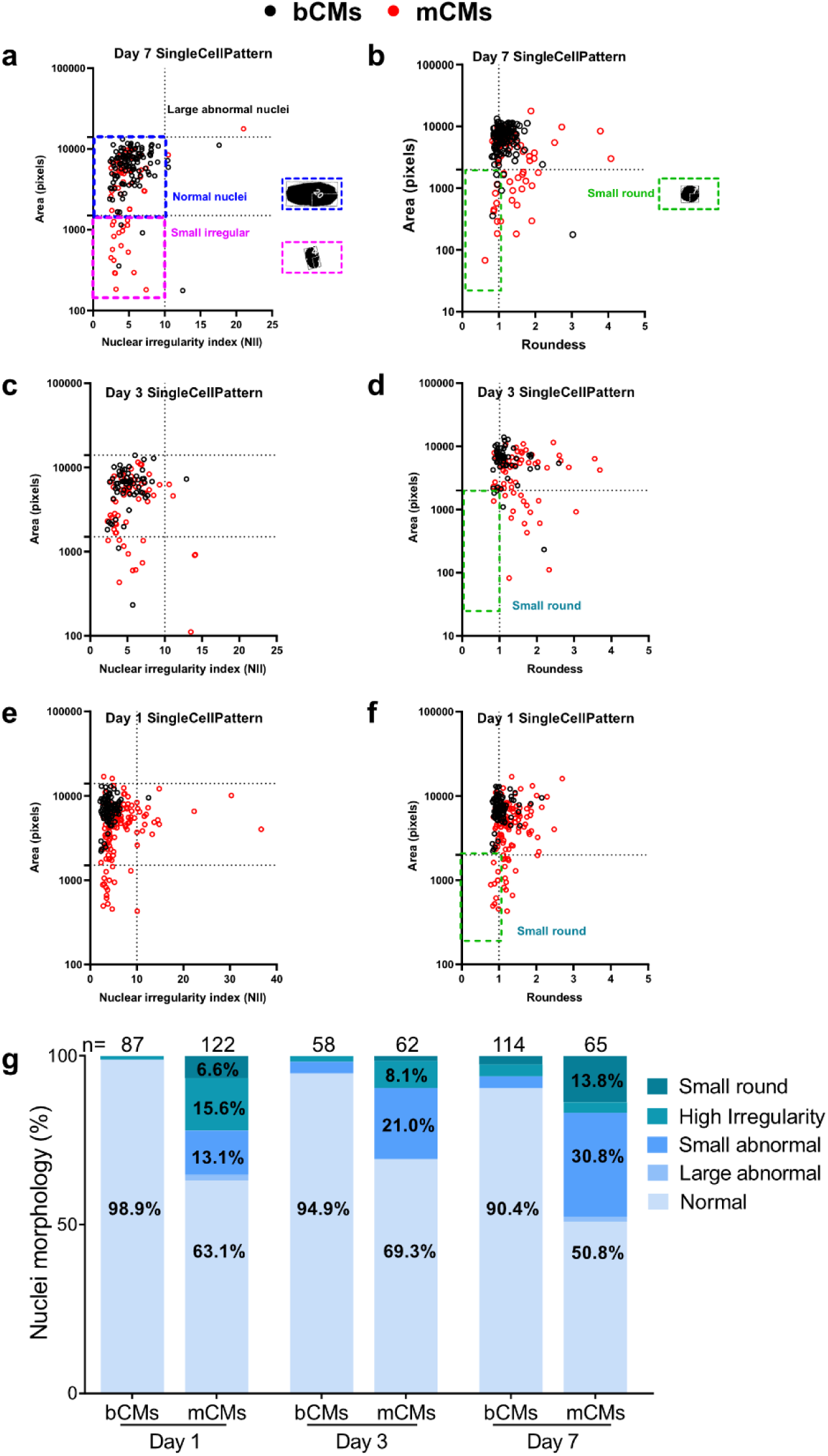
Nuclear morphology of micropatterned bCMs and mCMs. bCMs and mCMs were recovered from cryo and plated on single cell ECM rectangles. Unbiased analysis of nuclear morphology was performed using the Nuclear Morphometric Analysis plugin for ImageJ at day 7 **(a-b)**, 3 **(c-d)** and 1 **(e-f)**. Blue boxes indicated nuclei with normal morphology, violet boxes nuclei with small irregular morphology and green boxes nuclei with small round morphology characteristic of apoptosis. Representative examples are shown to the right of plots a-b. **(g)** Quantification of normal and abnormal nuclei from day 1 to 7. Chi-square test: Day 1, p<0.0001; Day 3, p<0.0001; Day 7, p<0.0001. Number of nuclei analyzed is indicated above bars. Bioreactor-derived cardiomyocytes, bCMs; Monolayer-derived cardiomyocytes, mCMs.

**Fig. S6:**
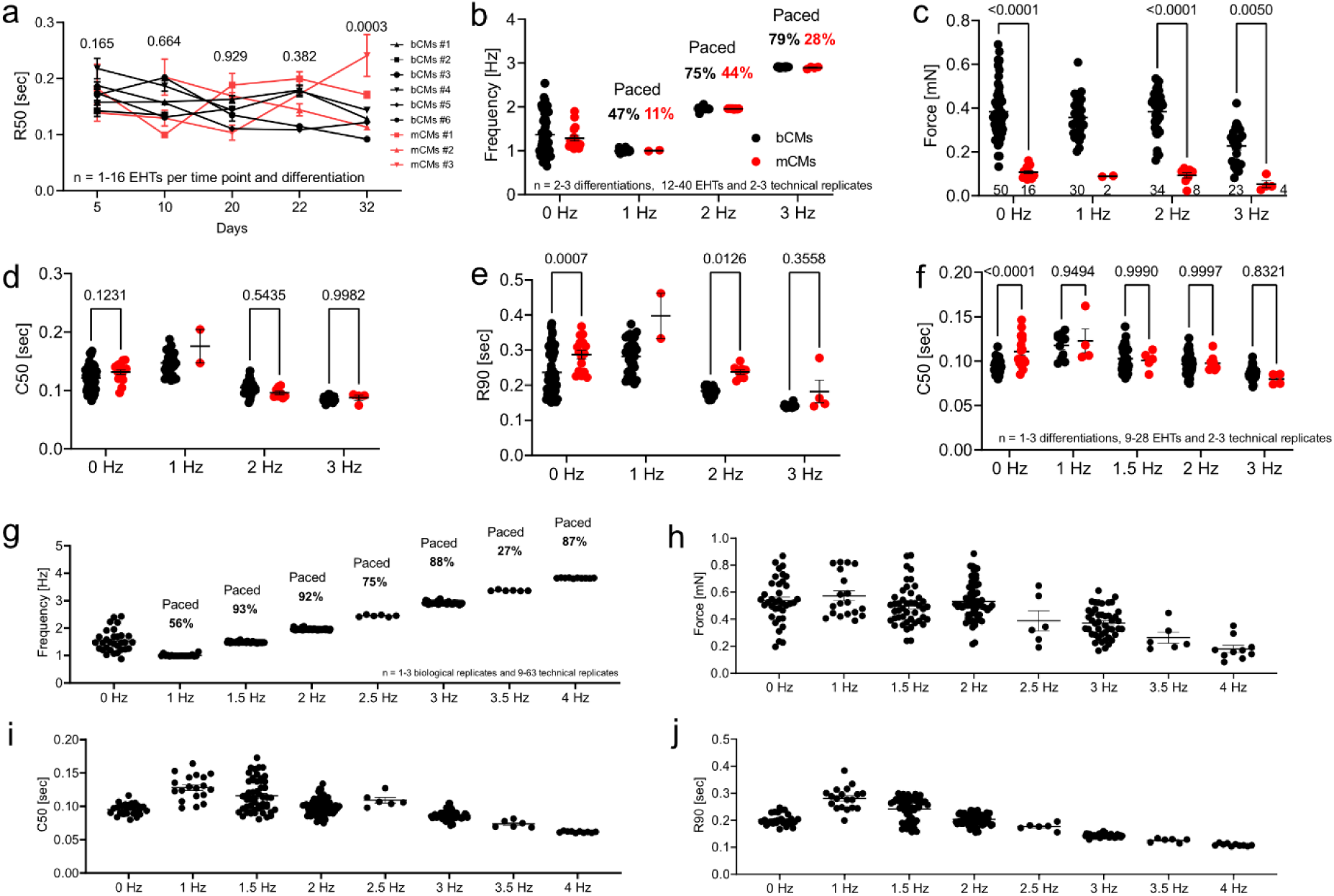
Characterization of EHTs assembled from bCMs or mCMs. Spontaneously beating EHTs assembled from cryo-recovered bCMs or mCMs were recorded in culture medium at 37°C from day 5 to day 32. Analyses of baseline 50% relaxation (R50; **a**). **(b-f)** Analysis of EHTs in culture medium (b-e) or Tyrode solution (f) without pacing (0 Hz) or with 1-3 Hz pacing. **b**, EHT beat frequency in response to pacing. Only EHTs captured by pacing are shown. The percent of EHTs captured at each pacing rate is indicated. Two-way ANOVA with Sidak’s post-test. **(g-h)** Extended graphs of Fig. 4f-h and Fig. S6f showing frequency **(g)**, force **(h)**, C50 **(i)** and R90 **(j)** measurements of bCM EHTs in Tyrode solution at 37°C and paced at 1, 1.5, 2, 2.5, 3, 3.5 and 4 Hz. Data are expressed as mean ± SEM. Bioreactor-derived cardiomyocytes, bCMs; Monolayer-derived cardiomyocytes, mCMs.

### Tables

All tables will be available in the revised manuscript.

**Table S1. Prior 2D and 3D protocols for iPSC differentiation to iPSC-CMs.** The table summarizes key features of selected iPSC-CM differentiation protocols and the properties of the resulting cells.

**Table S2. Markers of cell clusters in bCM and mCM iPSC-CM scRNAseq data.** Marker genes of each cell cluster are shown.

**Table S3. Estimate of cost of materials for monolayer or bioreactor iPSC-CM differentiation.** Labor costs are excluded. Monolayer costs are for 125 million cells. 100 ml bioreactor culture, or 125M monolayer cells. Yield is about 0.28 M/ml mCM vs 1.23 M/ml bCM. Yield per input iPSC is about 1.7 (mCM) vs 2.5 (bCM).

**Table S4. Antibodies used in this study.**

### Movie Legends

All movies will be available in the revised manuscript.

**Movie S1. Immunofluorescent analysis of a bioreactor-derived embryoid body at day 15 of differentiation.** Staining was performed against cTnT (red), ACTN2 (green) and nuclei (blue).

**Movie S2. Bioreactor-derived hiPSC-CMs at day 15 of differentiation. Movie S3. Monolayer-derived hiPSC-CMs at day 15 of differentiation.**

**Movie S4. Bioreactor-derived embryoid body beating at day 5 of differentiation.**

**Movie S5. Engineered heart tissue casted with cryopreserved bioreactor-derived hiPSC-CMs after 29 days in vitro.** Scale: 1 mm.

**Movie S6. Engineered heart tissue casted with cryopreserved monolayer-derived hiPSC-CMs after 29 days in vitro.** Scale: 1 mm.

**Movie S7. Engineered heart tissue casted with cryopreserved bioreactor-derived and transduced with adenovirus ChR2-YFP after 7 days in vitro.**

**Movie S8. Engineered heart tissue casted with cryopreserved bioreactor-derived optogenetically paced at 4 Hz after 41 days in vitro.**

## Notes

### Competing Interest Statement

The authors have declared no competing interest.

